# Control of Motor Landing and Processivity by the CAP-Gly Domain in the KIF13B Tail

**DOI:** 10.1101/2022.08.09.503328

**Authors:** Xiangyu Fan, Richard J. McKenney

## Abstract

Microtubules are a major component of the eukaryotic cytoskeleton that play crucial roles in diverse cellular process. Posttranslational modifications (PTMs) of tubulin dimers regulate the dynamics and organization of microtubules, as well as the interactions between microtubules and microtubule-associated proteins (MAPs). One unique PTM that occurs on microtubules is the cyclical removal and re-addition of the C-terminal tyrosine of α-tubulin. CAP-Gly (cytoskeleton-associated protein glycine-rich) domain containing proteins specifically recognize tyrosinated microtubules, a property exploited to regulate and spatially localize diverse microtubule effectors. KIF13B is a member of the long-distance transport kinesin-3 family, and the only kinesin motor that contains a conserved C-terminal CAP-Gly domain. What role the CAP-Gly domain plays in KIF13B’s motility along microtubules is unknown. Here, we investigated the interaction between KIF13B’s CAP-Gly domain, and the tyrosinated C-terminal tail domain of α-tubulin. We found that KIF13B’s CAP-Gly domain strongly influences the initial motor-microtubule interaction, as well as the processive motility of KIF13B along microtubules. The effect of the CAP-Gly domain on kinesin-microtubule binding is specific to the nucleotide state of the motor domain, suggesting an interplay between the N-terminal motor domain and C-terminal CAP-Gly domain underlies the KIF13B-microtubule interaction. These results reveal that specialized kinesin tail domains play active roles in the initiation and continuation of motor movement.

## Introduction

Microtubules are a critical component of the eukaryotic cytoskeleton, and are essential for various cellular functions, such as cell division and motility, intracellular signaling and transport, cell differentiation, and generation of specific or-ganelles^1^. microtubules are polymers composed of tubulin dimers, themselves composed of various α- and β-tubulin isotypes^2^. In addition to tubulin isotypes, the cytoplasmic surface of the microtubule polymer is modified with a variety of post-translational modifications (PTMs) to the C-terminal tail domains of the tubulin dimers. The large variety of tubulin isotypes and PTMs is collectively referred to as the “tubulin code”^1, 3, 4^. Cells express enzyme systems that act as “writers” of the tubulin code and modify the surface of the microtubule in a spatiotemporally controlled manner^4, 5^. In addition, a large repertoire of microtubule-associated proteins (MAPs) in cells, including molecular motors, act as “readers” of the tubulin code. One example is the direct effect of tubulin polyglutamylation on the recruitment and severing activities of spastin and katanin^6–8^. While it is hypothesized that the intracellular localization and activities of molecular motors could be directly modulated by tubulin PTMs, abundant evidence for this idea remains elusive because of the complex environment of the cytoplasm and the relatively weak effects of tubulin PTMs on motor activity *in vitro*^9–11^.

One unique PTM on microtubules is the enzyme-driven tubulin tyrosination/detyrosination cycle, in which the encoded C-terminal tyrosine residue of mammalian α-tubulin can be removed from the polypeptide chain by the tubulin carboxypeptidase vasohibins (VASHs) with the cooperation of its cofactor, small vasohibin binding protein (SVBP)^12, 13^. The tyrosine can also be added back onto α-tubulin by a tubulin tyrosine ligase (TTL)^14, 15^. In most cell types, tyrosinated microtubules make up the bulk of the polymer mass, and are typically more dynamic than detyrosinated microtubules^16–18^. Tyrosinated and detyrosinated microtubules compose distinct subsets of microtubules that orient in opposite directions within neuronal compartments^19^. Detyrosinated microtubules are more stable than tyrosinated microtubules, and cells polarize detyrosinated microtubules towards the direction of cell migration^20^, suggesting distinct functionalities for the two populations of modified microtubule polymer. Thus, the tyrosination/detyrosination cycle specifies subpopulations of microtubules for specific cellular functions, presumably in large part through the effect of the modification on microtubule effector molecules.

Some MAPs contain an evolutionary conserved CAP-Gly domain that specifically recognizes tyrosinated microtubules^21^. Structural work has revealed that the CAP-Gly domain interacts with the C-terminal EEY/F sequence motifs of α-tubulin^22, 23^, or a highly similar region found in the microtubule end-binding (EB) protein family and Cytoplasmic Linker Protein (CLIP) family^22, 24–26^. Some CAP-Gly domain-containing proteins, such as CLIP-170 and CLIP-115, participate in the regulation of microtubule dynamics and the recruitment of proteins to the microtubule plus-end^27–30^. Tubulin tyrosination can also affect the targeting and movement of motor proteins along microtubules. The p150^Glued^ (hereafter p150) subunit of the dynactin complex also contains a CAP-Gly domain that is important for targeting the dynactin complex to microtubule plus-ends in cells and initiating retrograde dynein movement from these regions^31, 32^. *In vitro*, the p150 CAP-Gly domain strongly biases the landing of assembled dynein-dynactin-cargo adapter complexes onto tyrosinated, versus detyrosinated microtubules^33, 34^, but it is not required for sustained processive movement of these complexes after initiation of movement^33^. The p150 CAP-Gly domain also enhances the dynein-dynactin interaction with microtubules under load^35, 36^. Therefore, the p150 CAP-Gly domain directly affects the landing and force production properties of the cytoplasmic dynein-dynactin complex, perhaps representing largest effect of a tubulin PTM on molecular motor output measured to date.

Within the kinesin family, the microtubule depolymerizing activity of kinesin-13 (MCAK) is directly regulated by tubulin tyrosination^10, 37^. Tyrosination also affects the velocity and processivity of kinesin-2 family motors^10^. Interestingly, within the large kinesin-3 family, the KIF13B motor contains a predicted CAP-Gly domain located at the C-terminus of the motor. This domain is highly conserved in all organisms with an identified KIF13B homologue (Fig. S1). KIF13B is involved in various cellular processes, such as axon formation, neuronal cell polarity, regulation of angiogenesis, directional migration of the primordial germ cell in *Xenopus* embryos, the uptake of lipoproteins via endocytosis, and the transport of Rab6 secretory vesicles from the Golgi^38–49^. More recently, KIF13B has also been observed to move bi-directionally within mammalian primary cilia^50^. The role of KIF13B’s CAP-Gly domain in these processes is not clear, but deletion of the domain in mice results in mislocalization of the truncated motor and KIF13B cargo within cells^43^. Here, we characterized the role of CAP-Gly domain in KIF13B during its processive motion along microtubules using *in vitro* reconstitution with single molecule imaging. We found that the CAP-Gly domain directly affects KIF13B’s motility by increasing the landing rate and extending the run length of KIF13B along tyrosinated microtubules. Our findings reveal that specialized tail domains of kinesin motors play active roles in the initiation and continuation of motor movement during cargo transport.

## Results

### Characterization of Recombinant KIF13B

We expressed and purified full-length and truncated versions of the human KIF13B motor using baculovirus expression in insect cells (Fig. 1A). All constructs were purified via a C-terminal superfolder-GFP (sfGFP)-strepII tag and gel filtration to near homogeneity (Fig. 1B). In addition to the wild-type full-length motor which is expected to exist in an autoinhibited state^51^, we generated two single point mutants, KIF13B^V178Q^ and KIF13B^K414A^, that have been previously shown disrupt the autoinhibition mechanism of KIF13B^52^. Additionally, we generated constructs lacking either the conserved kinesin motor domain (KIF13^Δmotor^), or a large portion of the C-terminal tail domain (KIF13^Δtail^, Fig. 1A, B). Previous studies have found that tail-truncated KIF13B motors are monomeric and must dimerize to induce processive motion along microtubules^52, 53^, although inclusion of longer segments of the tail domain appear to result in an active dimeric motor^40, 49^. Therefore, the oligomeric state of the full-length KIF13B motor remains unclear. We utilized mass photometry^54^ to examine the oligomeric state of our purified motor preparations across a range of nanomolar concentrations relevant to our single molecule assays. We observed that the WT KIF13B motor was predominantly monomeric across a 20-fold concentration range, but we could detect a minor proportion of dimeric motors at the intermediate and high concentrations examined (Fig. 1A-C). The V178Q and K414A mutations have previously been suggested to disrupt the intramolecular autoinhibition of KIF13B, which altered the monomer-dimer equilibrium for tail-truncated KIF13B^52^. Consistently, we observed our full-length KIF13B^V178Q^ or KIF13B^K414A^ motor preparations contained a larger proportion of dimeric motors across all concentrations tested as compared to the WT construct (Fig. 1C). Therefore, we conclude that full-length KIF13B exists predominantly as a monomeric motor in solution and has a weak tendency to dimerize at higher concentrations. Dimerization of the motor is enhanced by disruption of the autoinhibition mechanism that operates through the interaction between motor domain and first coiled-coil region of the motor^52^.

**Figure 1:**
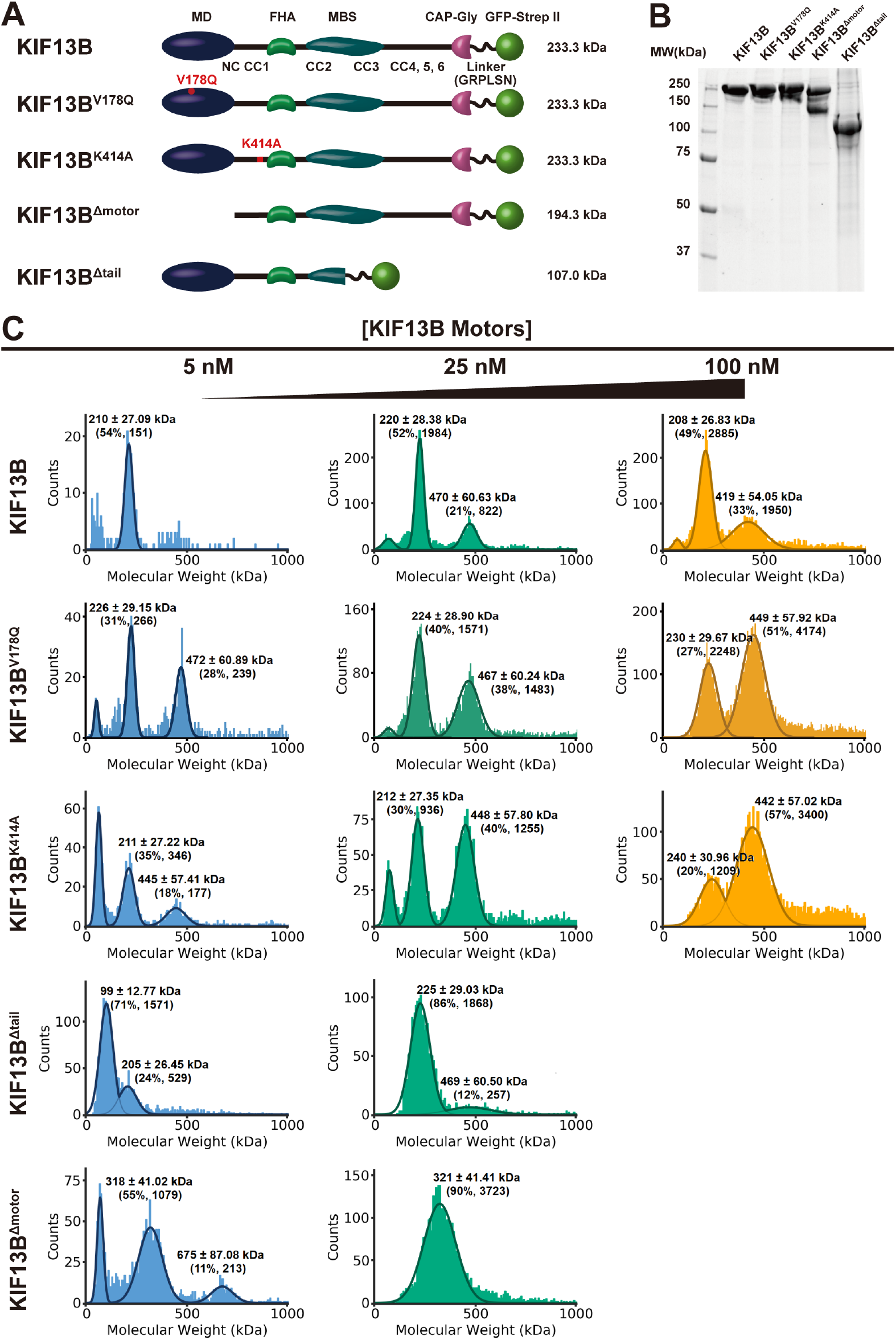
Mass photometry analysis reveals the oligomeric states of recombinant KIF13B motors. **A** Schematic of the KIF13B constructs used in this study. MD: motor domain, FHA: forkhead-associated domain, MBS: MAGUK (Membrane associated guanylate kinase) binding stalk, CC: coiled-coil, CAP-Gly: Cy-toskeleton-associated protein glycine-rich domain. Theoretical molecular weights of KIF13B motors are show to the right. **B** SDS-PAGE analysis of purified KIF13B motors. **C** Characterization of the oligomeric state of recombinant KIF13B motors by mass photometry. Histograms in each panel show the particle counts of recombinant KIF13B motors at the indicated molecular mass. For each protein, the molecular weights were measured at two or three different concentrations, from left panel to right panel: 5 nM, 25 nM and 100 nM. The dark lines are Gaussian fits to the peaks. The peak mass of the Gaussian fit, percentage of particles in the peak and the count of particles are indicated above each peak. The small peak at ~50 kDa mass in full length motors may represent proteolytic products around this size that were also observed in SDS-gels in Fig. 1B. For each concentration of recombinant proteins, the measurement was performed three times, and repeated with two different protein preparations.

We also examined the oligomeric state of our truncated motor constructs. We truncated KIF13B after the predicted second coiled-coil, and within the Membrane associated guanylate kinase (MAGUK) binding stalk (MBS) domain^55^ to generate KIF13B^Δtail^ (Fig. 1A-C). At lower concentrations, KIF13B^Δtail^, existed predominantly as a monomer with a smaller population of dimeric motors. However, raising the concentration only 5-fold resulted in near complete dimerization of this construct, revealing that removal of the C-terminal tail domain of KIF13B facilitates a much more dramatic concentration dependent dimerization of the motor (Fig. 1A-C), as compared to the full-length protein. For reasons that remain unclear, deletion of the kinesin motor domain resulted in a tail construct, KIF13B^Δmotor^, that had a mass most close to a trimeric state (Fig. 1C) at all concentrations examined. It is possible that removal of the kinesin motor domain results in aberrant oligomerization of the tail domain. The measured mass corresponds to a dimer of the full-length protein and a single copy of the major truncation product observed in the prep (Fig. 1A), which co-segregates with the construct through gel filtration. We were unable to record data for either truncated construct at concentrations higher than 25 nM due to overly high binding density of these proteins to the coverslip surface. From these data, we conclude that the C-terminal region of KIF13B negatively impacts the dimerization of the motor, suggesting that molecular interactions within this region could modulate the oligomeric state of the motor *in vivo*. A very recent structure of an autoinhibited and monomeric kinesin-3 member, KLP-6, reveals extensive interdomain interactions within the C-terminal tail of this kinesin-3 family member supporting this idea^56^.

### KIF13B binds preferentially to Tyrosinated microtubules

Because the tyrosination state of the microtubule lattice has been shown to affect the interactions of various motor proteins with microtubules^10, 19, 33, 57^, we sought to investigate whether the KIF13B CAP-Gly domain plays any direct role in the motor’s interaction with the microtubule lattice. In principle, KIF13B has two distinct types of microtubule-binding domains, the N-terminal ATP-dependent kinesin motor domain, and the C-terminal CAP-Gly domain (Fig. 1A, 2A).

**Figure 2:**
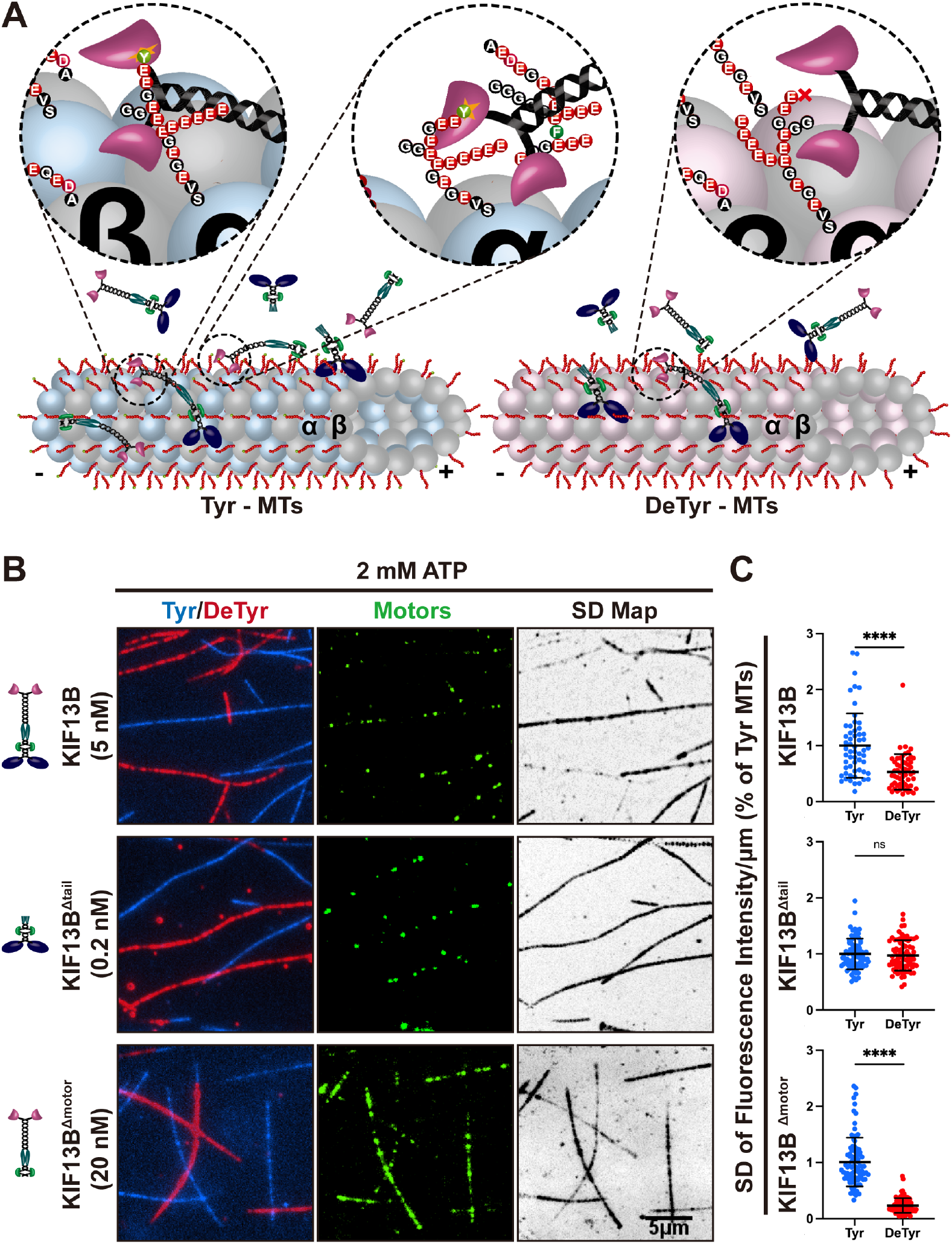
KIF13B prefer bind to tyrosinated microtubule. **A** Schematic showing the interaction between CAP-Gly domain of KIF13B with the tyrosinated tail of α-tubulin within microtubules. Left: On tyrosinated microtubules, the CAP-Gly domain of KIF13B and KIF13B^Δmotor^ can interact with the tyrosinated tail of α-tubulin. Right: On detyrosinated microtubules, this interaction is abolished. Circled regions showing high magnification views of the interactions between CAP-Gly domain of KIF13B motors with tyrosinated tail of α-tubulin. Tyrosinated and detyrosinated α-tubulin are colored in light blue and light pink respectively. **B** TIRF images of KIF13B motors (green, middle panel) bound to either tyrosinated (blue, left panel) or detyrosinated microtubules (red, left panel) in the presence of ATP. SD map: standard deviation maps from an entire time sequence reveal ensemble binding and dissociation events that lead to variations in pixel intensity, highlighting differences in binding between tyrosinated and detyrosinated microtubules. Scale bar: 5 μm. **C** Quantification of mean intensity (arbitrary units) from the SD maps per μm microtubules for recombinant KIF13B motors bound to the tyrosinated or detyrosinated microtubules relative to tyrosinated microtubules in the same chamber. KIF13B: n = 57, KIF13B^Δtail^: n = 80, KIF13B^Δmotor^: n = 89, microtubules quantified for each condition from two independent experiments. **** P< 0.0001. Mean ±SD are shown.

The relative role of each of these domains in the overall motor-microtubule interaction is unknown. To examine this, we directly compared the behavior of our KIF13B constructs on tyrosinated versus detyrosinated microtubules *in vitro* (Fig. 2A-B). In this assay, microtubules were polymerized from purified porcine brain tubulin, which is approximately 50% tyrosinated^58^. Carboxypeptidase A (CPA) treatment was used to remove the C-terminal tyrosine of α-tubulin^16, 33^. CPA treatment strongly reduced the binding of p150 to microtubules (Fig. S2A) as previously reported^33^, confirming detyrosination of the lattice. Differentially labeled WT (tyrosinated) and CPA-treated (detyrosinated) microtubules were mixed together in the same imaging chamber (Fig. 2B), providing an equal opportunity for motors to interact with each type of lattice.

Fluorescent proteins inter-acting transiently with the mi-crotubule surface generate large pixel intensity variations over background which can be averaged over an entire time series to generate a pixel-level fluorescence standard deviation (SD) map of the entire image series^33, 59^. The SD map for full-length KIF13B revealed a robust ~2-fold preference for binding to tyrosinated versus detyrosinated microtubules in the presence of ATP (Fig. 2B, C). In contrast, KIF13B^Δtail^, which lacks the CAP-Gly domain, displayed no preferential binding to either type of microtubule lattice (Fig. 2B, C), revealing the KIF13B motor domain displays no preference for the tyrosination state of the lattice. Removal of the kinesin motor domain resulted in an even stronger ~ 5-fold enhancement of binding on tyrosinated versus detyrosinated microtubules for KIF13^Δmotor^, suggesting the truncation binds microtubules predominantly via the CAP-Gly domain interaction with the tyrosinated α-tubulin tail domain (Fig. 2B, C). Residual binding of KIF13B^Δmotor^ to detyrosinated microtubules may be due to incomplete removal of the C-terminal tyrosine on the α-tubulin C-terminal tail (CTT). We conclude that KIF13B’s CAP-Gly plays a major role in the motor-microtubule interaction by strongly biasing the motor towards tyrosinated versus detyrosinated microtubules. Because the KIF13B motor domain is not affected by the tyrosination state of the lattice (Fig. 2B), these results reveal that the CAP-Gly domain enhances the initial encounter rate or enhances the interaction time of the motor on tyrosinated microtubules, or both.

### KIF13B’s CAP-Gly Domain Regulates the Motor’s Landing Rate and Run-length

We sought to detail how the CAP-Gly domain of KIF13B affects the motile parameters of KIF13B during the initiation and continuation of processive movement along microtubules. We utilized single-molecule microscopy to track the behavior of full-length KIF13B, its two active mutants KIF13B^V178Q^ and KIF13B^K414A52^ (Fig. 1A), along with the tail-truncated KIF13B^Δtail^. Purified motors were introduced into flow chambers containing differentially labeled tyrosinated and detyrosinated microtubules at low enough concentrations to discern individual motors and their behavior over time in the presence of ATP. We observed processive motility along microtubules from all of the motor constructs examined (Fig. 3A). The landing rate of KIF13B was more than 50-fold less than that of truncated or mutant motors (Fig. 3B), reflecting the strong autoinhibition of the wild-type motor^52^, and its predominantly monomeric state (Fig. 1C). We noticed that all full-length KIF13B motors showed fewer processive events on detyrosinated versus tyrosinated microtubules, while the motility of KIF13B^Δtail^ was largely similar between the two lattices (Fig. 3A, C). Indeed, quantification of the landing rates (all observable interactions of motors with the lattice) of each motor on each type of microtubule lattice revealed distinct differences between full-length motors containing the CAP-Gly domain, and KIF13B^Δtail^. Full-length motors had approximately two-fold higher landing rates on tyrosinated versus detyrosinated microtubules (Fig. 3B), consistent with our fluorescence intensity analysis (Fig. 2B,C). In contrast, we observed no change in landing rates between the different types of microtubules for KIF13B^Δtail^, revealing the preference for tyrosinated microtubules is driven by the tail domain of KIF13B. Importantly, we observed very similar re-sults with detyrosinated microtubules generated *in vitro* via CPA treatment, or treatment with the recently identified cellular tubulin detyrosinating enzyme complex, VASH1-SVBP^12, 13^ (Fig. S2), confirming the effect is directly related to the tyrosination status of the microtubule lattice.

**Figure 3:**
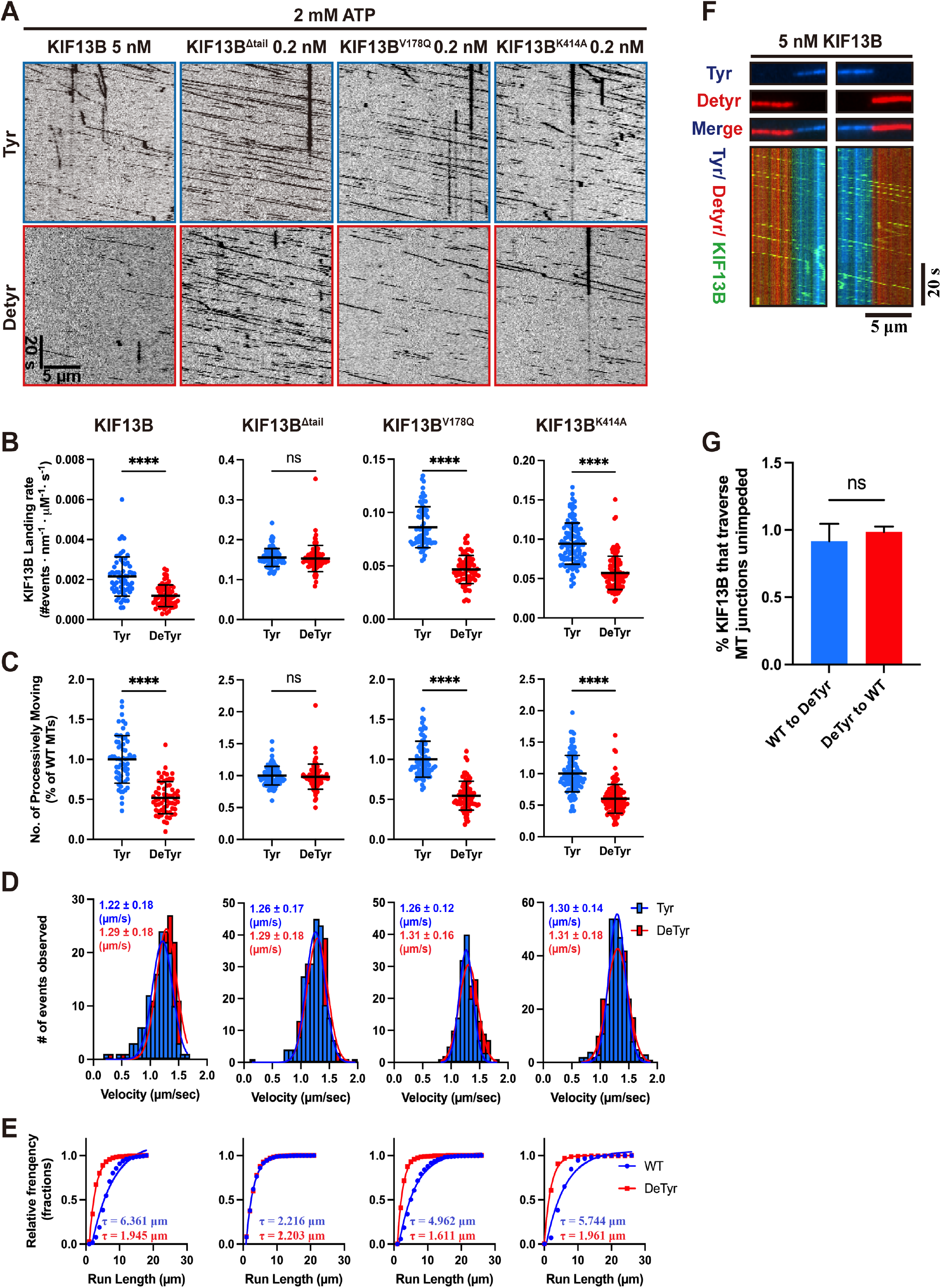
Regulation of motility by the CAP-Gly Domain. **A** Representative kymographs showing motility of KIF13B motors on tyrosinated (top panel) or detyrosinated microtubules (bottom panel) in the presence of 2 mM ATP. Scale bars: 20 s and 5 μm. **B** Quantification of the landing rates of KIF13B motors on tyrosinated and detyrosinated microtubules. For quantification in both B and C, microtubules were quantified for each condition from two independent experiments. KIF13B: n = 61 (both tyrosinated and detyrosinated microtubules). KIF13B^Δtail^: n = 81. KIF13B^V178Q^: n = 78. KIF13B^K414A^: n = 110). **** P< 0.0001. Mean ±SD are shown. **C** Quantification of the number of processive motors of KIF13B and its mutants per μm microtubule per second on tyrosinated or detyrosinated microtubules relative to tyrosinated microtubules in the same chamber. **** P< 0.0001. Mean ±SD are shown. **D** Quantification of the velocities of KIF13B motors on tyrosinated and detyrosinated microtubules. Histograms of the velocities were plotted for each population of motors and fit to a single Gaussian. The peak represents the average velocity of KIF13B or its mutants. Events were quantified for each condition from two independent experiments. KIF13B: n = 140, 152 (on tyrosinated and detyrosinated microtubules respectively). KIF13B^Δtail^: n = 232, 234. KIF13B^V178Q^: n = 152, 169. KIF13B^K414A^: n = 281,264. The mean ± SD for KIF13B and each mutant of KIF13B are indicated in the top left corners. **E** Cumulative frequency of the run lengths is plotted for each population of motors and fit to a one-phase exponential decay function. For quantification, events were quantified for each condition from two independent experiments. The characteristic run lengths derived from the fits (τ) are indicated. KIF13B: n = 414, 583 and R^2^= 0.958, 0.995 (on tyrosinated and detyrosinated microtubules respectively). KIF13B^Δtail^: n = 1004, 933 and R^2^= 0.999, 0.997. KIF13B^V178Q^: n = 517, 488 and R^2^= 0.985, 0.997. KIF13B^K414A^: n = 506, 528 and R^2^= 0.958, 0.997. **F** TIRF images (top) and kymographs (bottom) from chimeric microtubules composed of tyrosinated and detyrosinated microtubules. Motors moving either from tyrosinated onto detyrosinated microtubules (left panel in bottom) or from detyrosinated onto tyrosinated microtubules (right panel in bottom) are shown. Scale bars: 20 s and 5 *μ*m. **G** Quantification of the number of processive motors of KIF13B that traverse the microtubule boundaries and continue moving unimpeded. Motors were quantified for each condition from two independent experiments, n=77, 39 for motors moving from tyrosinated to detyrosinated microtubules and from detyrosinated to tyrosinated microtubules respectively. Mean ±SD are shown.

We next quantified KIF13B’s motile parameters on tyrosinated and detyrosinated microtubules located in the same chamber. All KIF13B constructs moved along both types of microtubules with a similar velocity of ~1.25 μm/s, consistent with a previous study^53^, revealing that the rate of the mechanochemical cycle of KIF13B is not affected by the tyrosination state of the microtubule (Fig. 3D). We next measured motor run lengths on each type of microtubule. Full-length KIF13B motors showed long continuous runs (Fig. 3A) with a fitted run length of ~ 5-6 μm on tyrosinated microtubules (Fig. 3E). Strikingly, motor run-lengths were reduced approximately two to three-fold on detyrosinated microtubules for all full-length motors examined (Fig. 3E), suggesting that an interaction of the tail domain with the tyrosinated microtubule lattice directly impacts motor processivity. In contrast, KIF13B^Δtail^ had approximately two to three-fold shorter runs than full-length motors (Fig. 3E), which we hypothesize is possibly due to weaker dimerization of the motor at the low concentrations used for single molecule assays (Fig. 1C). However, we note that the run lengths for the full length KIF13B are much longer than KIF13B^Δtail^ on tyrosinated microtubules (Fig. 3E), even though full length KIF13B is also predominantly monomeric at low concentrations (Fig. 1C). Therefore, it is possible that the full tail domain is required for stable dimeric assembly of KIF13B during processive motion, and more work, including structural analysis, is needed to fully address this question.

Importantly, no measurable difference in run-length was observed for KIF13B^Δtail^ on the two types of microtubule lattice (Fig. 3E). Additionally, we again found very similar effects on the motile parameters of full-length motors on detyrosinated microtubules generated by either CPA or VASH1-SVBP, confirming the specificity of CPA treatment (Fig. S3). Based on these data, we conclude that the CAP-Gly domain of KIF13B enhances both the motor’s landing rate, and processive run lengths, on tyrosinated microtubule lattices. In the absence of the CAP-Gly domain, the motor domain of KIF13B behaves indistinguishably on tyrosinated and detyrosinated microtubules lattices. Therefore, the tail domain of KIF13B, not the motor domain, recognizes the tyrosination state of the lattice and directly impacts the motility of the motor domain.

These results raised the possibility that the C-terminal CAP-Gly domain is continually engaged with the microtubule lattice during processive motility. Indeed, CAP-Gly domain containing proteins, such as p150, have been observed to diffuse, along the microtubule lattice^60^, providing a possible mechanism for interaction with the lattice during motility. To investigate this idea, we generated single chimeric microtubules composed of differentially labeled tyrosinated and detyrosinated sections of lattice^33^ (Fig. 3F). Using this system, we could observe how individual KIF13B motors behaved as they traversed from one type of lattice onto the other. We surmised that the CAP-Gly domain would be able to remain engaged with the tyrosinated, but not the detyrosinated lattice during motility. To our surprise, the majority of motors did not show obvious changes in the motile behavior as they traversed from one type of lattice to the other, but instead continued processive motility unabated (Fig. 3F). We quantified the fraction of motors that smoothly traversed the junction between each type of lattice and found that ~ 90% continued their processive motility while changing onto a different type of lattice (Fig. 3G). Because the CAP-Gly domain binds only weakly to detyrosinated lattices (Fig. 2B), we conclude that processive KIF13B motility does not require the CAP-Gly engagement with the microtubule lattice. However, the strong effect of detyrosination on motor run lengths (Fig. 3E) leads us to suggest that the CAP-Gly domain may allow for rapid motor re-attachment to the tyrosinated microtubule lattice upon dissociation of the motor domains during a processive run, facilitating continuation of processive motility.

### KIF13B’s Preference for Tyrosinated Microtubules is Nucleotide Dependent

Having observed that the KIF13B CAP-Gly domain regulates the motor’s landing rate and processive motility along microtubules, we next sought to investigate the molecular mechanism behind these observations. KIF13B contains two distinct types of microtubule binding domains. The kinesin motor domain cycles between a weak microtubule binding state (ADP state) and a strong microtubule binding state (ATP and apo states)^61^. In contrast, the non-enzymatic CAP-Gly domain only depends on the tyrosination state of the lattice for its interaction with microtubules. We sought to understand the relative contributions of each domain within the context of the KIF13B mechanochemical cycle. We performed microtubule binding assays with tyrosinated and detyrosinated microtubules in the same chamber, and KIF13B motors in either saturating ADP to lock the motor domain in the weak binding state, or the ATP analogue Adenylyl-imidodiphosphate (AMP-PNP) to lock the motor domain in the strong binding state (Fig. 4A). In the presence of saturating ADP, we observed that full length KIF13B binding to tyrosinated microtubules was ~8-fold higher than on detyrosinated microtubules in the same chamber (Fig. 4B), suggesting that when the motor domain is bound to ADP, KIF13B’s interaction with the microtubule is strongly dominated by the CAP-Gly domain. In contrast, KIF13B binding to tyrosinated microtubules was only ~ 1.4-fold higher than to detyrosinated microtubules in AMP-PNP (Fig. 4B), suggesting that in the strong binding state of the ATPase cycle, the affinity of the motor domain for the lattice largely overwhelms the preference of the CAP-Gly domain for tyrosinated microtubules. We performed the same experiment with the tail-truncated construct, KIF13B^Δtail^, and observed no difference in binding to tyrosinated or detyrosinated microtubules in the presence of ADP or AMP-PNP (Fig. 4B), consistent with our prior results that the motor domain of KIF13B is not affected by the tyrosination state of the lattice (Fig. 3A, B).

**Figure 4:**
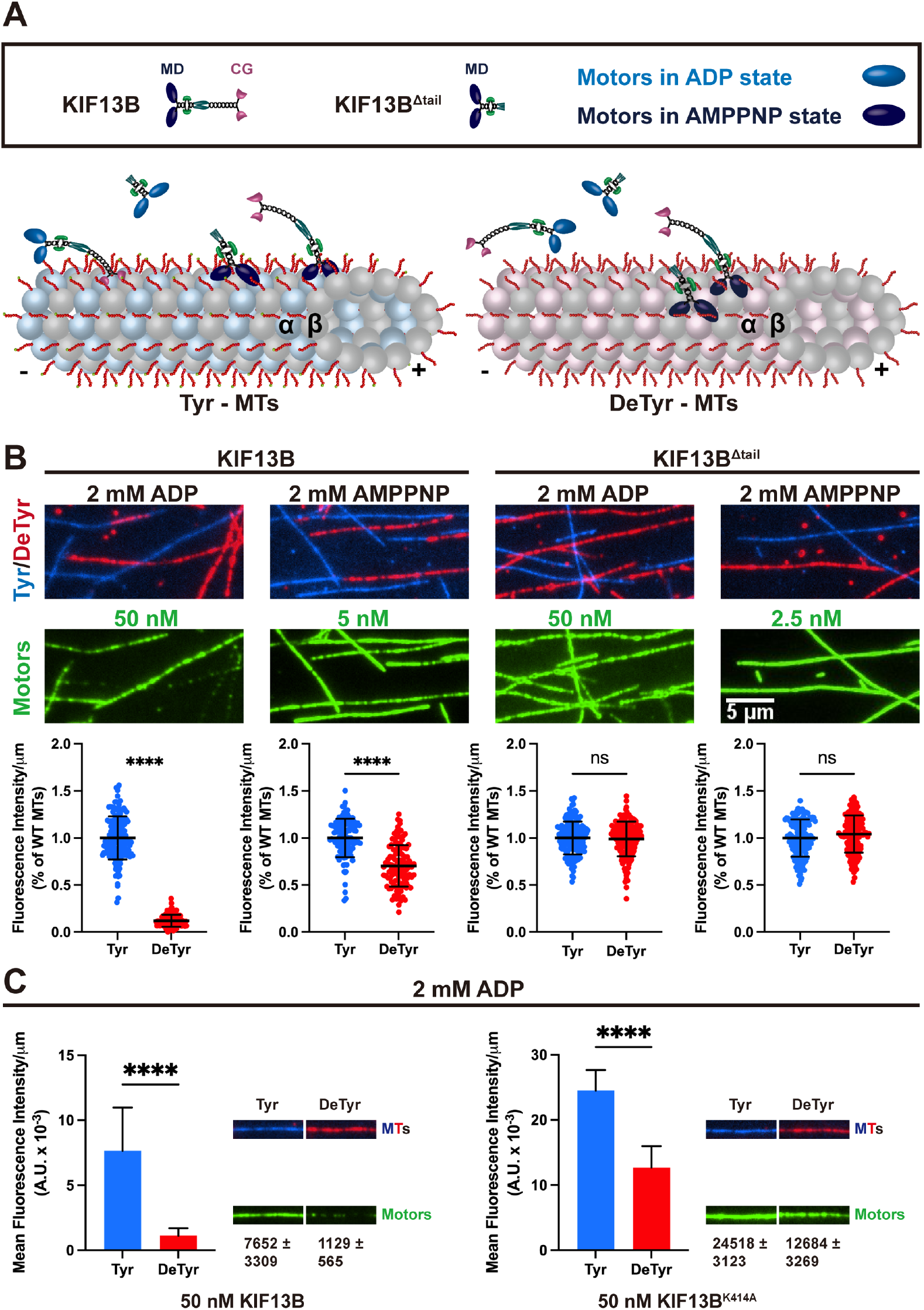
Nucleotide dependent preference for tyrosinated microtubules of KIF13B. **A** Schematic showing interactions between KIF13B motors and tyrosinated or detyrosinated microtubules in the presence of 2 mM ADP or 2 mM AMPPNP. Left: motors binding along the tyrosinated microtubules. In ADP sate (motor domains: light blue), the CAP-Gly domain dominates the interaction with microtubules. In the AMPPNP state (motors domains: dark blue), the motor domain dominates the interaction with microtubules. Right: motors binding along the detyrosinated microtubules. Since the interaction between CAP-Gly and α-tubulin is abolished, the motor domain dominates the interaction between microtubules and motors in either ADP or AMPPNP states. MD: Motor Domain, CG: CAP-Gly Domain. **B** The fluorescence intensity of KIF13B and KIF13B^Δtail^ on tyrosinated and detyrosinated microtubules in different nucleotide sates. TIRF images of KIF13B and KIF13B^Δtail^ (green, middle panel) bound to either tyrosinated (blue, top panel) or detyrosinated microtubules (red, top panel). Bottom panel, quantification of mean fluorescence intensity (arbitrary units) per μm microtubules for motors bound to tyrosinated or detyrosinated microtubules relative to tyrosinated microtubules in the same chamber. Microtubules were quantified for each condition from two independent experiments. KIF13B (in AMPPNP state): n = 116 (both tyrosinated and detyrosinated microtubules). KIF13B^Δtail^ (in AMPPNP state): n = 150. KIF13B (in ADP state): n = 164. KIF13B^Δtail^ (in ADP state): n = 160. Scale bar: 5 *μ*m. **** P< 0.0001. Mean ±SD are shown. **C** Comparison of the microtubule affinities of KIF13B motors on tyrosinated and detyrosinated microtubules in the presence of 2 mM ADP. 50 nM of KIF13B or KIF13B^K414A^ was incubated with tyrosinated and detyrosinated microtubules in chambers for 10 min, mean fluorescence intensities of the MT-bound KIF13B (left) or KIF13B^K414A^ (right) were quantified (left side bar graphs). Representative microtubule images and the corresponding motor channels are shown in right side in each condition. The width of each panel corresponds to 8.06 μm. Mean fluorescence intensity of motors ± SD is shown below. For quantification, microtubules were quantified for each condition from two independent experiments. For each condition, n = 80. **** P< 0.0001.

To understand how autoinhibition and dimerization affects the ability of the CAP-Gly domain to direct KIF13B to tyrosinated microtubules, we directly compared the fluorescence intensity of KIF13B and KIF13B^K414A^ in the presence of ADP. We assayed the motors at 50 nM, a concentration at which we expected the fraction of dimeric KIF13B^K414A^ motors to be substantially higher than that for KIF13B (Fig. 1C). Comparison of the average motor density along microtubules revealed substantially higher levels of KIF13B^K414A^ bound to both tyrosinated and detyrosinated microtubules as compared to KIF13B (~ 3-fold and 11-fold higher for KIF13B^K414A^ on tyrosinated and detyrosinated microtubules respectively, Fig. 4C). Importantly, an ~ 2-fold preference of KIF13B^K414A^ for tyrosinated over detyrosinated microtubules remained, consistent with the idea that the CAP-Gly domain directs the motor’s interaction with microtubules when the motor domain is bound to ADP. We conclude that release from autoinhibition and dimerization of the motor results in a higher affinity for both types of microtubules. Because the amount of KIF13B^K414A^ bound to detyrosinated microtubules in the presence of ADP is much higher than KIF13B, we hypothesize that that avidity arising from dimerization of the motor enhances the binding affinity of both the CAP-Gly and motor domains within KIF13B. In addition to potential avidity effects, the increased affinity is also likely due to microtubule-stimulated release of ADP from the motor domain, which is enhanced upon release from the autoinhibited state^52^. From these data, we conclude that the KIF13B’s preference for tyrosinated microtubules is nucleotide dependent. In the ADP state, the CAP-Gly domain dominates the interaction between KIF13B and microtubules, resulting in a prominent preference for tyrosinated microtubules. In the ATP state (mimicked by AMP-PNP), the motor domain dominates the interaction between KIF13B and microtubule, and the preference for tyrosinated microtubules is diminished. Because ADP release is the rate-limiting step in the kinesin mechanochemical cycle, and is greatly accelerated by microtubule binding^61^, the motor domain is expected to accumulate in this state prior to association with the microtubule, suggesting that the CAP-Gly domain will direct the initial encounter of KIF13B with microtubules in cells.

### KIF13B’s CAP-Gly Domain Enhances the Motor-Microtubule Interaction When the Motor Domain is in the ADP State

To further understand how the CAP-Gly domain affects KIF13B’s interaction with microtubules, we analyzed the microtubule binding behavior of KIF13B and its mutants on tyrosinated and detyrosinated microtubules in ADP state at the single molecule level. To visualize single motor binding events on microtubules, a relatively low concentration of motors was used, and based on our mass photometery results (Fig. 1C), the motors are likely predominantly monomeric. Kymographs showed that in the presence of ADP, single molecules of KIF13B and its mutants bound to both types of microtubules (Fig. 5A). Quantification of the landing rates of these motors revealed that both KIF13B and its active mutate KIF13B^K414A^ showed a strong preference for tyrosinated microtubules, as the landing rates of both motors on tyrosinated microtubules was ~2-fold higher than on detyrosinated microtubules (Fig. 5B). Consistent with the ideas that this preference is driven by the CAP-Gly domain, KIF13B^Δmotor^ displayed an increased ~ 3-fold preference for tyrosinated microtubules, while KIF13B^Δtail^ again showed no preference for either type of microtubule (Fig. 5B).

**Figure 5:**
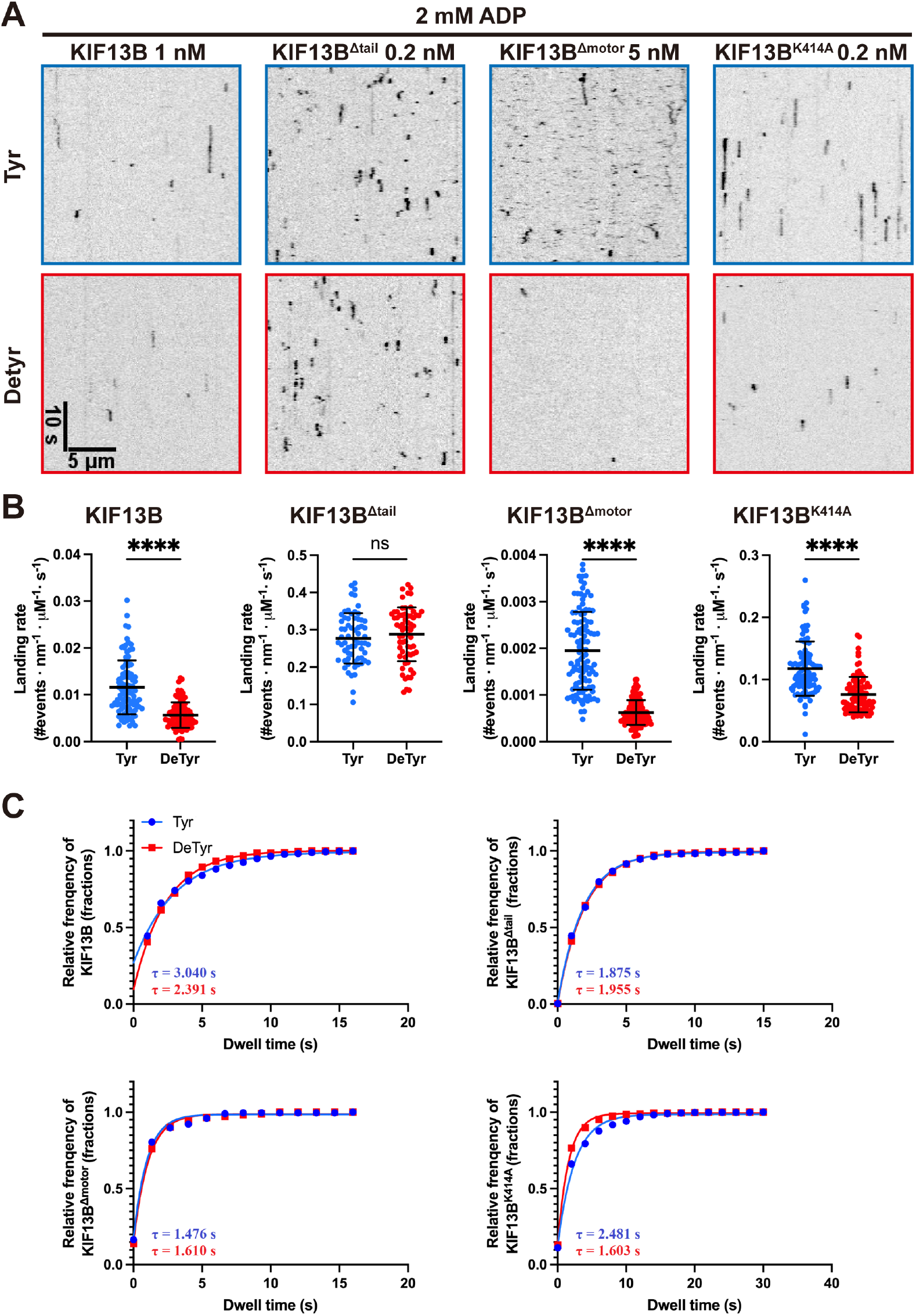
CAP-Gly domain enhances the interaction between KIF13B and tyrosinated microtubule in ADP state. **A** Representative kymographs showing KIF13B motors binding to tyrosinated (top panel) and detyrosinated microtubules (bottom panel) in the presence of 2 mM ADP. Scale bars: 10 s and 5 μm. **B** Quantification of landing rates of KIF13B motors on tyrosinated and detyrosinated microtubules in ADP state. For quantification, microtubules were quantified for each condition from two independent experiments. KIF13B: n = 96 (both on tyrosinated and detyrosinated microtubules). KIF13B^Δtail^: n = 65. KIF13B^Δmotor^: n = 112. KIF13B^K414A^: n = 94). **** P< 0.0001. Mean ±SD are shown. **C** Quantification of the dwell times of KIF13B and its mutants on tyrosinated or detyrosinated microtubules. Cumulative frequency of the dwell time is plotted for each population of motors and fit to a one-phase exponential decay function. The characteristic dwell times derived from the fits (τ) are indicated in the bottom left corners. For quantification, numerous of events dwell time longer than 3 pixels were selected and quantified, two independent experiments were performed for each condition. KIF13B: n = 414, 317 and R^2^= 0.989, 0.999 (on tyrosinated and detyrosinated microtubules respectively). KIF13B^Δtail^: n = 433, 470 and R^2^= 0.999, 0.9998. KIF13B^Δmotor^: n = 204, 285 and R^2^= 0.991,0.997. KIF13B^K414A^: n = 520, 482 and R^2^= 0.989, 0.997.

We next measured the dwell times of motors from the resulting kymographs. This analysis revealed that while KIF13B^Δtail^ showed highly similar dwell times on both types of microtubules, the other constructs tested showed longer dwell time events on tyrosinated microtubules (Fig. 5A-C). We found that the dwell times of KIF13B and KIF13B^K414A^ on tyrosinated microtubules were ~1.3 and ~1.5 times higher than on detyrosinated microtubules respectively, while KIF13B^Δmotor^ and KIF13B^Δtail^ showed no difference in dwell times (Fig. 5C). These results suggest that the CAP-Gly domain enhances the motor’s dwell time on tyrosinated microtubules when the motor domain is in the ADP state, revealing a possible mechanism for CAP-Gly dependent initiation of processive motility on tyrosinated microtubules.

### Model for the role of the CAP-Gly domain in KIF13B motility

Our data reveal at least two roles of the CAP-Gly domain in KIF13B motility. Firstly, when the motor domains are bound to ADP and not engaged with the microtubule lattice, the CAP-Gly domain facilitates the interaction of the motor with the microtubule through its binding of the α-tubulin C-terminal tyrosine residue (Fig. 6, step 1). The oligomerization state of the motor at this step is currently unclear, but we suggest that the binding of the CAP-Gly domain to the lattice could facilitate localized concentration of motors on the lattice, in turn stimulating dimerization and activation of processive motility. It is also conceivable that engagement of the CAP-Gly domain with the microtubule leads to molecular rearrangements of the motor from the autoinhibited, monomeric state, priming the motor for dimerization on the microtubule surface. A similar mechanism has recently been suggested for myosin-7a^62^. Alternatively, binding of cargo to the motor may stimulate dimerization prior to microtubule association. Association of the CAP-Gly with the microtubule could facilitate the binding of the motor domains to the lattice, leading to nucleotide exchange (Fig. 6, step 2), and subsequent processive motility (Fig. 6, step 3). During processive motility, the motor domains remain out of phase with respect to nucleotide state such that one motor domain is always tightly bound to the lattice. However, at some frequency, both motor domains enter the weak binding ADP state simultaneously, leading to release from the lattice. The CAP-Gly domain may facilitate fast rebinding of the motor to the lattice, preventing diffusion away from the microtubule and termination of motility (Figure 6, step 4).

**Figure 6:**
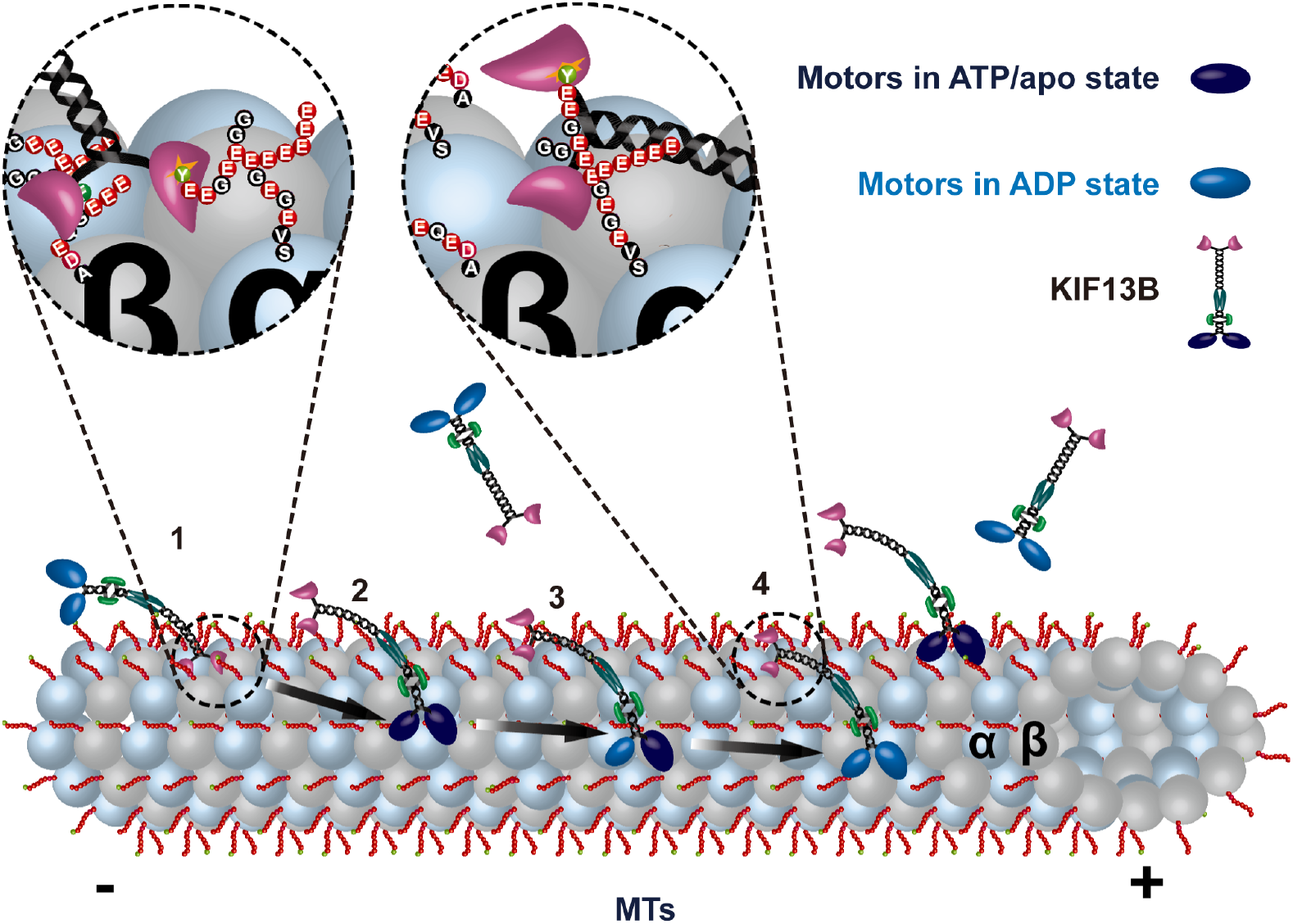
Model for the regulation of CAP-Gly domain in KIF13B motility. Schematic showing a model for the roles of the CAP-Gly domain in KIF13B motility: (1) In the ADP state, the CAP-Gly domain dominates the interaction between KIF13B and microtubules. Interaction between the CAP-Gly domain and the tyrosinated α-tubulin within microtubules facilitates the binding of the motor domain to microtubule lattice. (2) Landing of the motor domain onto the microtubule stimulates ADP releasing, leading to nucleotide exchange, which in turn increases affinity between KIF13B and microtubules. (3) Nucleotide exchange begins processive motility along microtubules powered by asynchronous ATP hydrolysis in the two motor domains. In this process, at least one motor domain remains strongly bound to the lattice until: (4) at some time frequency, both motor domains simultaneously enter the weak binding ADP state, leading to motor release from the lattice. In this case, the interaction between CAP-Gly domain and tyrosinated tail of α-tubulin facilitates rebinding of the motor to the lattice and initiates a new mechanochemical cycle.

## Discussion

Kinsein-3 motors transport cargos over long distances within cells and are critical for human health and disease^63^. As such, understanding the molecular mechanisms underpinning the functions of this kinesin family represents an important question in cell biology. Here we have studied the function of the unique tail domain of KIF13B, which contains a specialized second microtubule binding site in its CAP-Gly domain. Our work revealed that KIF13B’s CAP-Gly domain drives a strong preference for motor binding to tyrosinated MTs *in v*itro, and that this preference depends on the nucleotide state of the motor domain. In addition, the CAP-Gly domain mediates higher landing rates and an approximately 3-fold longer run length on tyrosinated versus detyrosinated MTs. Given that defects in kinesin-3’s interactions with microtubules underly human neurodevelopmental and neurodegenerative diseases^64, 65^, our results provide new insights into the various strategies adopted by kinesin-3 motors to augment the ability of the motor domains to move processively along microtubules to support cellular trafficking homeostasis and human health.

Previous studies have revealed that CAP-Gly domain containing proteins CLIP-170, CLIP-115 and their orthologues in yeast are involved in the regulation of microtubule dynamics, via their specific recruitment to the dynamic plus end of MTs^27–30^. The CAP-Gly domains of these proteins directly interact with the EEY/F motif of α-tubulin^22, 23^ and a similar motif found on end-binding (EB) proteins^29, 66^. The microtubule plus end localization of these proteins stimulates the dynamicity of MTs by increasing castastrophy and rescue frequencies^27–30^. Given that many CAP-Gly proteins are recruited to dynamic microtubule plus-ends through interactions with EB proteins, we note that despite a current lack of evidence, it is formally possible that KIF13B is also recruited to microtubule plus-ends in a similar manner. Human KIF13B contains several candidate SxIP motifs known to be crucial for interactions with EBs^67^, and future work should focus on a potential role for the plus-end recruitment of KIF13B in cellular biology.

In the context of motor protein movement, *in vitro* reconstitution showed that the CAP-Gly domain within the p150 subunit of the dynactin complex strongly modulates the initiation of dynein-driven motility along tyrosinated MTs without affecting the continuous movement of the DDB complex^33^, while the domain is also important for the initiation of retrograde movements in cells^34^. These studies illustrate that CAP-Gly domain mediated interactions are utilized by cells to fine tune both microtubule dynamics and the initiation of movement of motor proteins. We now extend these findings to the kinesin family of antero-grade motors and suggest that KIF13B cargo transport dynamics within cells is likely influenced by the tyrosination state of the local microtubule lattice.

KIF13B has been reported to transport various cargos within cells, such as: Phosphatidylinositol 3,4,5-triphosphate (PIP3) containing lipid vesicles, low-density lipoprotein (LDL) receptor-related protein 1 (LRP1) containing vesicles, Vascular endothelial growth factor receptor 2 (VEGFR2) and Myosin X^40–49^. The molecular basis of these diverse motor-cargo interactions remains unclear, but some interactions involve the forkhead-associated (FHA) domain or MBS regions of the motor^47, 55^ (Fig. 1A). To date, a direct role for the distal CAP-Gly domain in cargo-binding has not been described, but its deletion in mice results in reduced uptake of LRP1^43^. Our results imply that misdirection of the motor along the differentially tyrosinated microtubules may underlie this observation.

KIF13B is not the only kinesin that contains auxiliary microtubule binding domain. Kinesin-1 is also reported to contain a secondary microtubule-binding site composed of charged residues located within the C-terminal tail of the heavy chain, which is important for its ability to reorganize the cytoskeleton by sliding microtubules relative to one another^68, 69^. The highly processive motor kinesin-8 (mammalian KIF18A and yeast Kip3p) contains a microtubule-binding site in its tail domain is essential for correct plus end localization of this motor in mitosis^70^. Therefore, specialized microtubule binding sites located distal to the kinesin motor domain are an apparently malleable evolutionary advantage for diverse kinesin functions in cells and our results build on this repertoire within the kinesin family.

Finally, a recent structure of the autoinhibited kinesin-3 family member KLP-6 revealed a multimodal mechanism for keeping the monomeric motor in an inactive state^56^. The mechanism involves intricate folding of most of the kinesin tail domain around the motor domain, sterically occluding the microtubule binding site and trapping the motor in the ADP state. This mechanism likely has strong parallels in KIF13B, as the domain organization of these motors is largely similar. If the CAP-Gly domain of KIF13B plays a role in autoinhibition of the motor is an interesting question for future structural studies of KIF13B and its homologs. Our *in vitro* reconstitution with full-length KIF13B will provide a platform to examine new hypotheses about the mechanism and regulation of this important transport kinesin.

## Acknowledgements

We thank all the members of the MOM lab for their continual input and feedback on this project. The authors thank Shinsuke Niwa (Tohoku University, Japan) for generously providing the purified VASH1-SVBP complex. The work was supported by grants from NIGMS GM124889 (to RJM). This paper was typeset with the bioRxiv word template by @Chrelli: www.github.com/chrelli/bioRxiv-word-template

## Author contributions

R.J.M. and X.F. designed the research. R.J.M. secured research funding. X.F performed the research and analyzed the data. R.J.M. and X.F wrote the paper. All authors edited the paper.

## Materials and Methods

### Plasmids

Full-length human KIF13B cDNA was purchased form Transomics (BC172411). A DNA fragment encoding the sfGFP-2xStrepII tag was synthesized by gBlocks (Integrated DNA Technologies, Coralville, IA, USA). To generate insect cell expression constructs, Coding sequences of human KIF13B, KIF13B^Δmotor^ (a.a. 358-1826), KIF13B^Δtail^ (a.a. 1-679), KIF13B^V178^Q and KIF13B^K414A^ were tagged with sfGFP-2xStrepII in C-terminal and cloned into pAceBac1 vector (Geneva Biotech) by Gibson Assembly. Coding sequence of Full length KIF13B, KIF13B^Δtail^ and sfGFP-2xStrepII were amplified by PCR, all point mutations were introduced by PCR-based mutagenesis. All constructs were verified by Sanger sequencing.

### Protein expression and purification

For protein expression, baculovirus expression system was used. Briefly, insect Sf9 cells were grown or maintained in Sf-900 II serum-free medium (Thermo Fisher Scientific) or in ESF medium (Expression Systems) at 27°C. Plasmids DNA of corresponding constructs were transformed into DH10EmBacY competent cells (Geneva Biotech) to generate bacmids. To get baculovirus, 1 x 10^6^ Sf9 cells were transferred to each well of a six-wells plate and subsequently transfected with 1 μg of bacmid DNA by 6 μl of Cellfectin (Thermo Fisher Scientific). Baculovirus containing cell supernatants (P1) were harvested 7 days (cells in Sf-900 II medium) or 11 days (cells in ESF medium) after transfection when the entire cell population has become infected. To amplify the baculovirus (P2), 50 ml of Sf9 cells were infected with 50 μl of P1 virus, the P2 virus were harvested 4 days (cells in Sf-900 II medium) or 7 days (cells in ESF medium) after infection. To prepare recombinant proteins, 200 ml of Sf9 cells at high density (~2 × 10^6^cells/ml) were infected by P2 virus (virus: cells = 1:100, v/v) and cultured around 65 hrs. at 27°C. Cells were subsequently harvested by centrifuge at 1,000 x g for 10 min and flash frozen by liquid nitrogen. Frozen cells were stored at −80°C before purification.

For recombinant protein purification, frozen pellet was thawed on ice and resuspended in 40 ml of PB buffer (purification buffer: 50 mM Tris-HCl pH 8.0, 150 mM KCH_3_COO, 2 mM MgSO_4_, 1 mM EGTA, 5% glycerol) freshly supplemented with 1 mM dithiothreitol (DTT), 1 mM phenylmethylsulfonyl fluoride (PMSF), 1 mM ATP, and protease inhibitor mix (Promega). Resuspended cells were homogenized in a dounce homogenizer and lysed by addition of 1% Triton X-100 and 5 μl of Benzonase Nuclease (Millipore) for 10 min on ice. To obtain soluble lysate, lysed cells were centrifuged at 15, 000 × g for 20 min at 4°C. For affinity purification, the clarified lysate was incubated with 2 ml of Strep-Tactin XT resin (IBA) for 1 hr at 4°C and washed with PB buffer, protein was subsequently eluted by elution buffer (PB buffer plus 100 mM biotin, pH 8.0). All proteins were further purified by gel filtration. Briefly, eluted protein was concentrated to 500 μl via Amicon Ultra 5 centrifugal filters (Merck) and separated by Phenomenex Yarra 3 μm SEC-4000 300 × 7.8 mm column (Phenomenex) in GF150 buffer (25 mM HEPES (pH 7.4), 150 mM KCl, and 1 mM MgCl_2_) supplemented with 0.01% NP-40. Protein containing fractions were pooled and supplemented with 0.1 mM ATP and 10% Glycerol. Protein aliquots were subsequently flash frozen by liquid nitrogen and stored at -80°C. Protein concentrations were measured by NanoDrop One (Thermo Fisher Scientific) based on the absorbance of fluorophores.

### Single molecule mass photometry assay

The single molecule mass photometry assays were carried out as previous described^54, 62^. Briefly, to prepare chambers, microscope cover glasses (#1.5 24 × 50 mm, Deckgläser) were cleaned by 1 hour sonication in Milli-Q H_2_O, followed by another hour sonication in isopropanol, cover glasses then washed by Milli-Q H_2_O and dried by filtered air. CultureWell silicone gaskets (Grace Bio-Labs) were cut and washed by Milli-Q H_2_O, CultureWell silicone gaskets were then dried by filtered air and placed onto the freshly cleaned cover glasses providing four independent sample chambers. Samples and standard proteins were diluted in HP buffer (90 mM HEPES, 10 mM PIPES, 50 mM KCH_3_COO, 2 mM Mg(CH_3_COO)2, 1 mM EGTA, 10% glycerol, pH = 7.0). The HP buffer was freshly filtered by 0.22 μm filter before measurement. For calibration, standard proteins BSA (Sigma), Apoferritin (Sigma) and Thyoglobulin (Sigma) were diluted to 10-50 nM. The no-specific binding events of single molecules on cover glasses were scattered and the masses of these molecules were measured by OneMP instrument (Refeyn) at room temperature. Data were collected at an acquisition rate of 1 kHz for 100 s by AcquireMP (Refeyn) and subsequently analyzed by DiscoverMP (Refeyn).

### Microtubule preparation

Porcine brain tubulin was isolated using the high-molarity PIPES procedure and then labelled with biotin NHS ester, Dylight-405 NHS ester or Alexa647 NHS ester as described previously (https://mitchison.hms.harvard.edu/files/mitchisonlab/files/labeling_tubulin_and_quantifying_labeling_stoichiometry.pdf). Pig brains were obtained from a local abattoir and used within ~4 hrs. after death. To polymerize microtubules, 50 mM of unlabeled tubulin, 10 μM of biotin-labeled tubulin and 3.5 μM of Dylight-405-labeled (or Alexa647-labelled) tubulin were incubated with 2 mM of GTP for 20 min at 37°C. Polymerized microtubules were stabilized by addition of 20 μM taxol and incubated additional 20 min. microtubules were pelleted at 20, 000 × g by centrifugation over a 150 μl of 25% sucrose cushion and the pellet was resuspended in 50 μl BRB80 (80 mM Pipes (pH 6.8), 1 mM MgCl_2_ and 1 mM EGTA) containing 10 μM taxol.

Carboxypeptidase A (CPA)-treated detyrosinated microtubules were prepared as previous description^33^. Briefly, 50 mM of unlabeled tubulin, 10 μM of biotin-labeled tubulin and 3.5 μM of Alexa647-labelled microtubules tubulin were incubated with 2 mM of GTP and 12 mg/ml CPA (Sigma) for 20 min at 37°C, followed by the addition of 20 μM taxol for an additional 20 min. The digestion was stopped by the addition of 10 mM DTT, and the CPA enzyme was removed by centrifugation of the microtubules over a 25% sucrose cushion and the pellet was resuspended as described above.

To prepare chimeric microtubules, stabilized tyrosinated and detyrosinated microtubules were equally mixed and incubated overnight at room temperature for end-to end annealing. Annealed microtubules were verified by TIRF microscopy.

### Total Internal Reflection (TIRF) assays

TIRF chambers were assembled from acid washed glass coverslips as previously described (http://labs.bio.unc.edu/Salmon/protocolscoverslip-preps.html), pre-cleaned slide and double-sided sticky tape. Chambers were first incubated with 0.5 mg/ml PLL-PEG-biotin (Surface Solutions) for 5-10 min, followed by 0.5 mg/ml streptavidin for 5 min. Microtubules were diluted by BRB80 contain 10 μM taxol. Diluted microtubules were flowed into streptavidin adsorbed flow chambers and incubated for 5 to 10 min at room temperature for adhesion. To remove unbound microtubules, chambers were subsequently washed twice by HP assay buffer (90 mM HEPES, 10 mM PIPES, 50 mM KCH3COO, 2 mM Mg(CH3COO)2, 1 mM EGTA, 10% glycerol, bovine serum albumin (BSA) (1 mg/ml), biotin-BSA (0.05 mg/ml), K-casein (0.2 mg/ml), 0.5% Pluronic F-127 and 10 mM taxol, pH = 7.0). Purified motor protein was diluted to indicated concentrations in the assay buffer with 2 mM of corresponding nucleotides (ATP, ADP or AMPPNP) and an oxygen scavenging system composed of PCA/PCD/Trolox. Then, the solution was flowed into the glass chamber. Images were acquired using a Micromanager software-controlled Images were acquired using o Micromanager software-controlled Nikon TE microscope (1.49 numerical aperture, 100× objective) equipped with a TIRF illuminator and Andor iXon charge-coupled device electron-multiplying camera. Data were analyzed manually using ImageJ (Fiji), and statistical tests were performed in GraphPad Prism 9.

### Protein sequences alignment

Protein sequences of KIF13B homologs were retrieved from diverse eukaryotic organisms using Uniprot Knowledgebase (UniprotKB), and subsequently were aligned using Clustal WS with default settings of Jalview (Clustal W and Clustal X version 2.0)^71^.

### Preparation of VASH1-SVBP treated microtubules

To prepare VASH1-SVBP treated detyrosinated microtubules, 10 nM of purified VASH1-SVBP complex were added into polymerized microtubules above (Alexa647-labelled) and freshly supplemented with 1 mM of PMSF and 1 mM of DTT. The mixture was incubated overnight at 37°C to remove the tyrosine at C-terminal of α-tubulin in microtubules. The reaction was terminated by the addition of 10 mM DTT, and the VASH1-SVBP complex was removed by centrifugation of the detyrosinated microtubules at 20, 000 x g over a 25% sucrose cushion and the pellet was resuspended as described above.

## Supplementary Materials

**Figure S1:**
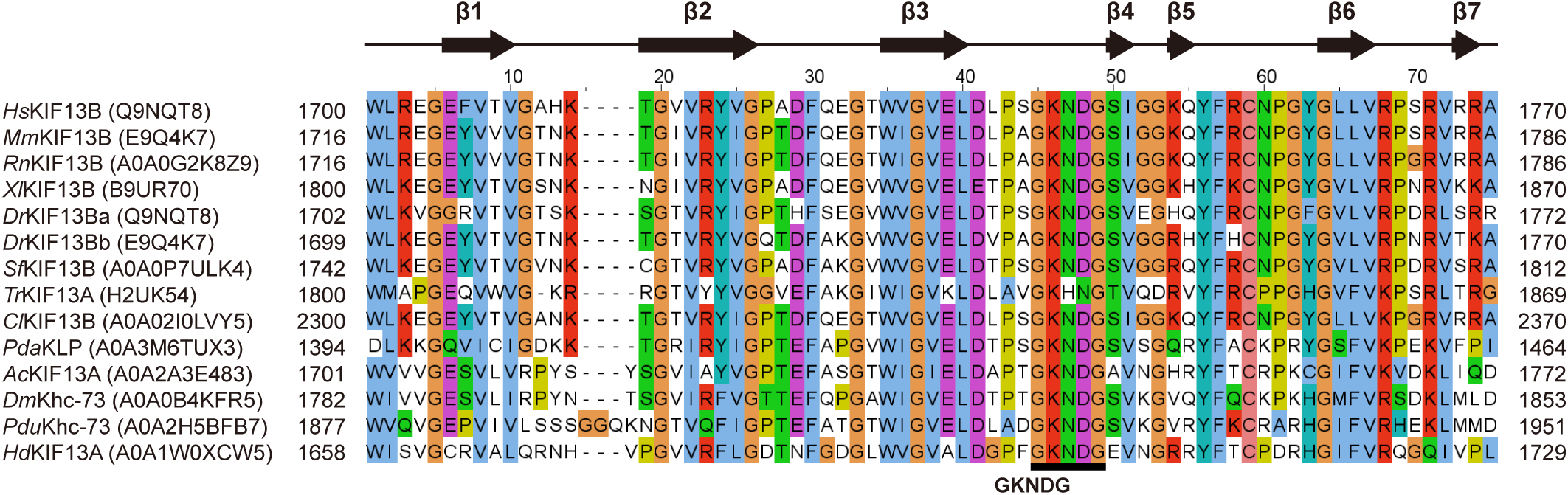
The CAP-Gly domain of KIF13B is evolutionary conserved. Alignment of CAP-Gly domain protein sequences in KIF13B homologues. The predicted secondary structure is shown above, arrows and lines representing β-strand and random coils respectively. The core GKNDG motif within CAP-Gly domain was underlined by black^22^. The abbreviations of species names and protein names, as well as UniProtKB accession numbers (in brackets) are indicated in the left side before the protein sequences. Numbers show the positions of residues in corresponding protein sequences. Species names and abbreviations: *Homo sapiens (Hs), Mus musculus (Mm), Rattus norvegicus (Rn), Xenopus laevis (Xl), Danio rerio (Dr), Scle-ropages formosus (Sf), Takifugu rubripes (Tr), Columba livia (Cl), Pocillopora damicornis (Pda), Apis cerana (Ac), Drosophila melanogaster (Dm), Platynereis dumerilii (Pdu), Hypsibius dujardini (Hd)*.

**Figure S2:**
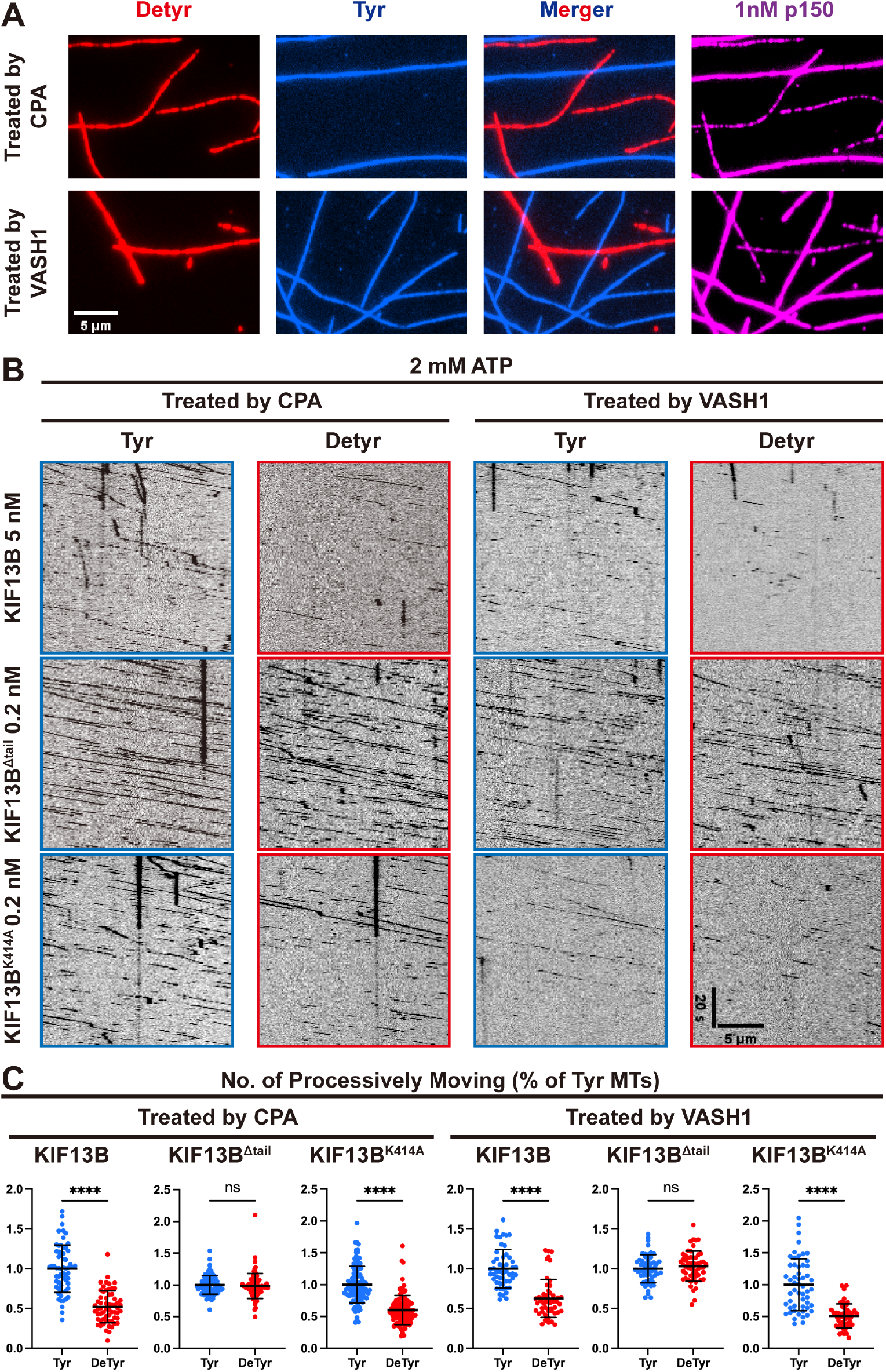
Quantification of recombinant KIF13B motors on detyrosinated microtubules treated by CPA or VASH1-SVBP. **A** CPA-treated (top panel, red) or VASH1-SVBP treated (bottom panel, red) microtubules were mixed with untreated microtubules (blue) and introduced into same chamber. As a control, 1 nM of p150 (magenta) was added and incubated 10 min at room temperature and subsequently visualized by TIRF microscopy. Scale bar: 5 μm. **B** Representative kymographs showing movements of KIF13B motors on tyrosinated (top panel) or detyrosinated microtubules (bottom panel). Left two panels, detyrosinated microtubule treated by CPA (reproduced from figure 2B for comparison). Right two panels, detyrosinated microtubule treated by VASH1-SVBP. Scale bars: 20s and 5 μm. **C** Quantification of the number of processive motors of KIF13B motors per μm MT per second on tyrosinated or detyrosinated microtubules relative to tyrosinated microtubules in the same chamber. Top panels, detyrosinated microtubule treated by CPA (data were re-plotted from figure 2D for comparison). Bottom panels, detyrosinated microtubule treated by VASH1-SVBP. For the quantification of microtubules treated by VASH1-SVBP, microtubules were quantified for each condition from two independent experiments. KIF13B: n = 52 (both tyrosinated and detyrosinated microtubules). KIF13B^Δtail^: n = 56 (both tyrosinated and detyrosinated microtubules). KIF13B^K414A^: n = 54, 51 (both tyrosinated and detyrosinated microtubules respectively). **** P< 0.0001. Mean ±SD are shown.

**Figure S3:**
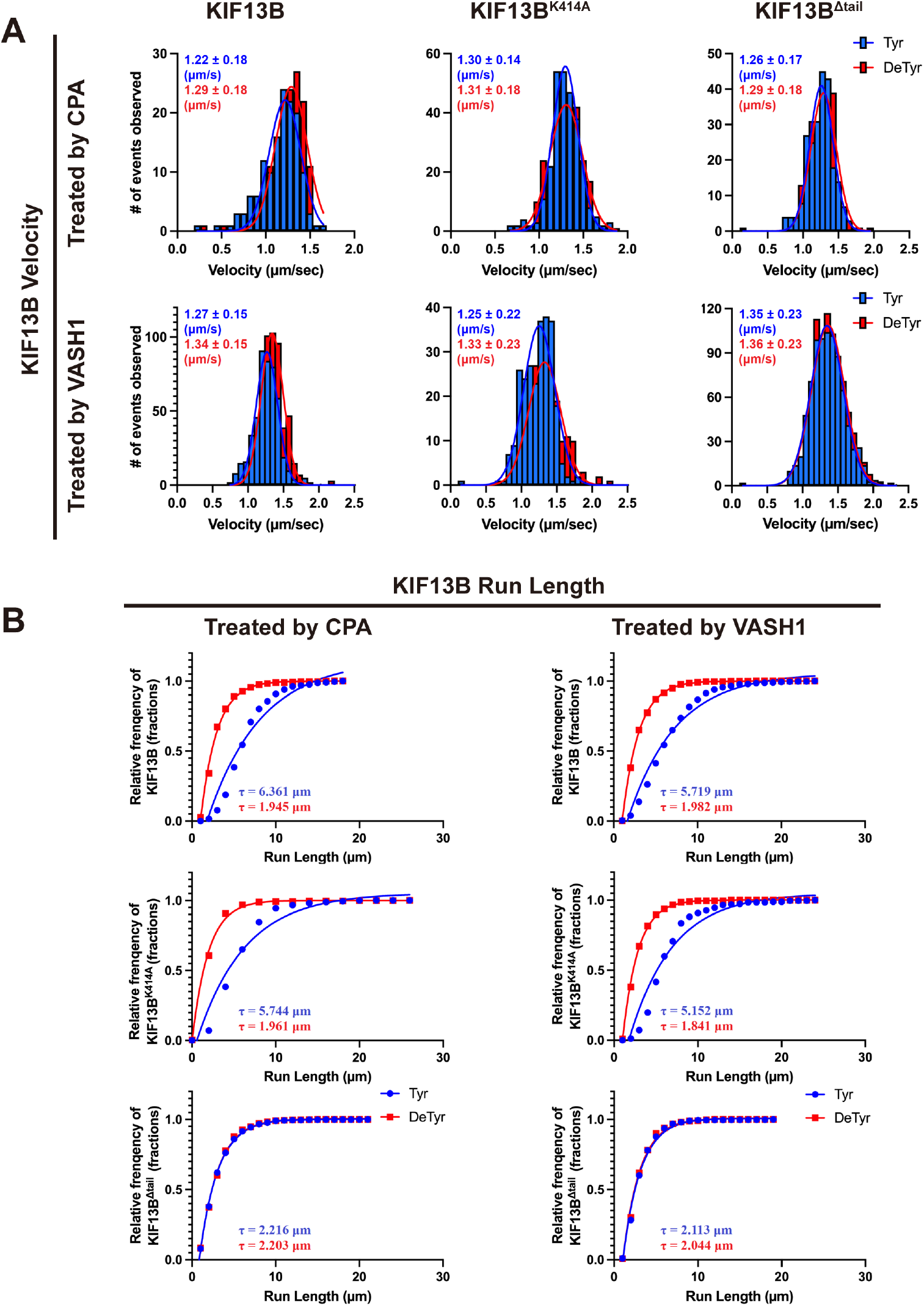
Comparison the motile parameters of recombinant KIF13B motors on detyrosinated microtubules treated by CPA or VASH1-SVBP. **A** Quantification of the velocities of KIF13B motors on tyrosinated and detyrosinated microtubules. Histograms of the velocities were plotted for each population of motors and fit to a single Gaussian. The peak represents the average velocity of KIF13B and corresponding mutants. Top panels, detyrosinated microtubule treated by CPA (data were re-plotted from figure 2E for comparison). Bottom panels, detyrosinated microtubule treated by VASH1-SVBP. For the quantification of motors on VASH1-SVBP treated microtubules, events were quantified for each condition from two independent experiments. The mean ± SD for wild-type and each mutant of KIF13B are indicated in the top left corners. KIF13B: n = 476, 564 (on tyrosinated and detyrosinated microtubules respectively). KIF13B^Δtail^: n = 838, 854. KIF13B^K414A^: n = 262, 213. **B** Quantification of the run length of KIF13B motors on tyrosinated and detyrosinated microtubules. Cumulative frequency of the run lengths is plotted for each population of motors and fit to a one-phase exponential decay function. Left panels, detyrosinated microtubule treated by CPA (data were re-plotted from figure 2F for comparison). Right panels, detyrosinated microtubule treated by VASH1. For the quantification of motors on VASH1-treated microtubules, numerous of events were quantified for each condition from two independent experiments. The characteristic run lengths derived from the fits (τ) are indicated in the bottoms. KIF13B: n = 476, 564 and R^2^= 0.978, 0.9995 (on tyrosinated and detyrosinated microtubules respectively). KIF13B^Δtail^: n = 838, 854 and R^2^= 0.992, 0.993. KIF13B^K414A^: n = 262, 213 and R^2^ = 0.956, 0.998. Mean ±SD are shown.

